# Effects of TMS on the decoding and electrophysiology of priority in working memory

**DOI:** 10.1101/2025.09.15.676374

**Authors:** Jacqueline M. Fulvio, Bradley R. Postle

**Affiliations:** Department of Psychology, University of Wisconsin–Madison; Department of Psychiatry, University of Wisconsin–Madison

## Abstract

The flexible control of working memory (WM) requires prioritizing immediately task-relevant information while maintaining information with potential future relevance in a deprioritized state. Using double-serial retrocuing (DSR) with simultaneous EEG recording, we investigated how single pulses of transcranial magnetic stimulation (spTMS) to right intraparietal sulcus impacts neural representations of unprioritized memory items (UMI), relative to irrelevant memory items (IMI) that are no longer needed for the trial. Twelve human participants (8 female) performed DSR plus a single-retrocue task, while spTMS was delivered during delay periods. Multivariate pattern analysis revealed that spTMS restored decodability of the UMI concurrent with stimulation, and that of the IMI several timesteps later, after the evoked effects of spTMS were no longer present in the EEG signal. This effect was carried by the alpha (8-13 Hz) and low-beta (13-20 Hz) frequency bands. Analyses of the raw EEG signal showed two effects selective to the epoch containing the UMI: the retrocue and spTMS each produced phase shifts in the low-beta band. These findings demonstrate that deprioritization involves active neural mechanisms distinct from the processing of the IMI, and that these are supported by low-beta oscillatory dynamics in parietal cortex. We hypothesize that the mechanism underlying spTMS-triggered involuntary retrieval of the UMI is the disruption of the encoding of priority status, which may depend on oscillatory dynamics in the low-beta band.

**Significance Statement:** This study provides key insights into how the brain maintains information at different levels of priority in working memory. Importantly, it shows how deprioritizing information in working memory is different from simply “dropping” it. By combining transcranial magnetic stimulation with high temporal resolution measurement of neural signals (with EEG), we replicate previous findings that spTMS triggers the involuntary retrieval of previously unprioritized information and provide new insight into how it does this. Specifically, at the level of EEG dynamics, spTMS is shown to uniquely alter oscillatory dynamics in the low-beta band.

## Introduction

The flexible control of behavior requires controlling the contents of our working memory (WM) to prioritize the information that is critical for in-the-moment behavior, to maintain information with potential future relevance in a way that does not interfere with current behavior, and to remove information that is no longer useful. These processes have been operationalized with the double-serial retrocuing (DSR) procedure in which the one of two memory items not cued by a first prioritization cue temporarily acquires the status of ‘unprioritized memory item’ (UMI). Critically, in DSR, the UMI remains relevant because it might be cued later in the trial (Lewis-Peacock et al., 2012).

Initial neuroimaging studies of the DSR task reported that multivariate pattern analysis (MVPA) evidence for an active representation of the UMI dropped to baseline during the delay following the first retrocue (with fMRI, Lewis-Peacock et al., 2012; Rose et al., 2016; LaRocque et al., 2017; with EEG, LaRocque et al., 2014; Rose et al., 2016). Furthermore, when a single pulse of transcranial magnetic stimulation (spTMS) was applied during this time, decodability of the UMI was recovered (Rose et al., 2016), a pattern consistent with the idea that prioritization may be accomplished, in part, via a transformation of the UMI into an “activity silent” state (c.f., Stokes, 2015). More recently, however, several studies have identified evidence for an active representation of the UMI. Some of these have trained and tested the decoder on the UMI itself (e.g., Christophel et al., 2018; Paluch et al., 2025), and some have trained on the PMI and tested on the UMI (i.e., cross-label decoding). Interestingly, cross-label decoding has yielded evidence that the active representation of the UMI is “opposite” that of the PMI, manifested as below-baseline decoding of the UMI (van Loon et al., 2018), as larger multivariate representational dissimilarity between PMI and UMI within category than between category (van Loon et al., 2018), and as ‘flipped’ inverted encoding model (IEM) reconstructions of the UMI relative to the PMI (Wan et al., 2020; Yu et al., 2020) consistent with a common representational format for UMI and PMI. (But see Iamshchinina et al. (2021) for cross-label decoding results that do not suggest “opposite” coding of the UMI.)

Although demonstrations of successful decoding of the UMI from neural signals raise questions about the activity-silent hypothesis (c.f., Barbosa et al., 2021, Wolff et al., 2021; Kandemir et al., 2024), understanding the mechanisms underlying the effects of spTMS on working memory remains important because spTMS influences behavior, not just decoding. For example, during blocks of 1-item recall-of-location trials, spTMS delivered during the intertrial interval (ITI) influences the magnitude of the attractive serial bias exerted by the previous trial’s location on the subsequent trial’s recall (Barbosa et al., 2020). In the DSR task, spTMS affects behavior in priority-dependent ways consistent with findings using brief 20Hz TMS trains to posterior sensory cortex (Zokaei et al., 2014a; Zokaei et al., 2014b). This is demonstrated with a version of the DSR procedure in which the UMI is presented as the recognition probe for a subset of the probes that require a “nonmatch” response. spTMS increases the false-alarm rate (FAR) to these lures when delivered prior to the first of the two recognition probes, but not when delivered later in the trial, when the item not cued by the second retrocue has the status of “irrelevant memory item” (IMI; because it will not be tested; Rose et al., 2016, Fulvio & Postle 2020). This would be expected if spTMS had the effect of reactivating only items currently held in WM.

What might be the neural bases of the effect of spTMS on the UMI? Initial evidence points at an important role for oscillatory dynamics in the beta band. In one study, filtering the EEG signal into classical frequency bands suggested that the spTMS restoration of the decodability of the UMI is restricted to the beta band (Rose et al., 2016). In another, the decomposition of the EEG signal into a set of discrete coupled-oscillator components revealed that only components with peaks in the beta band, and that were prominent at posterior scalp locations, were sensitive to spTMS in a manner that related to its behavioral effects (Fulvio et al., 2024). These latter results suggested that spTMS may exert its effects on DSR performance by disrupting a beta band-encoded deprioritization mechanism, thereby reducing priority-based discriminability of the cued item (“prioritized memory item” (PMI)) relative to the UMI due to an increased level of interitem competition. In the present study, we recorded the EEG during performance of DSR and a closely matched single-retrocue (SR) task, to assess the role of low-frequency dynamics, particularly in the low-beta band (13-20 Hz), in priority-related effects of spTMS on WM performance.

## Methods

### Code accessibility

The code/software described in the paper is freely available online at [URL will be provided prior to publication].

### Participants

Human subjects were recruited at a location which will be identified if the article is published. Twelve neurologically healthy individuals (8 females, 18-28 years, *M* = 28.3 years, all right-handed), with no reported contraindications for magnetic resonance imaging (MRI) or transcranial magnetic stimulus (TMS) took part in all portions of the study. The sample size was chosen according to a power analysis based on Experiment 3 (n = 6) from Rose et al. (2016), which demonstrated that spTMS delivered to posterior parietal cortex produced a brief (single timepoint) reemergence of the UMI in EEG-based decoding. The classification of the UMI significantly differed from chance at the timepoint immediately following the spTMS pulse, (*t*(5) = 4.35, *p* < .005; Rose et al., 2016, Supplementary Materials, p. 13). From this result, we calculated Cohen’s *d*, yielding an effect size of *d* = 4.35 / √6 = 1.78. Using G*Power 3.1, we conducted the power analysis for a one-sample *t*-test (one-tailed, testing whether post-spTMS classification accuracy would exceed chance), with α = .05, power = .80, and *d* = 1.78. This analysis indicated a required sample size of n = 4 participants would be needed. However, because significant findings from small-sample studies often overestimate the true effect size (because only unusually large effects are likely to reach statistical significance when sample sizes are small (Button et al., 2013)), a sample size of n = 4 was almost certainly insufficient for a robust replication. To account for the likelihood that the effect may be smaller in a new study, we adopted a more conservative effect size estimate of *d* = 0.80, which represents a large but realistic effect. We repeated the power analysis using this effect size and this analysis indicated a required sample size of at least n = 10 participants would be needed.

All participants had normal or corrected-to-normal vision with contact lenses (eyeglasses were not compatible with TMS targeting apparatus), and all reported having normal color vision. The research was approved by the [Author University] Health Sciences Institutional Review Board. All participants gave written informed consent at the start of each session and received monetary compensation in exchange for participation.

### Experimental procedure

The study was carried out in four sessions, each on a separate day. The first session was an MRI scan that yielded the anatomical image used to guide spTMS during the ensuing three behavioral EEG sessions. During each behavioral session, participants performed alternating blocks of DSR and SR, with each session comprising eight 30-trial blocks. Block order was counterbalanced across participants and switched across sessions within participants.

### Behavioral tasks

The single retrocue (SR) task began with the simultaneous presentation of two sample items drawn from two of three categories (face, word, direction of dot motion), one above and one below central fixation (2 sec), followed by 5 sec of fixation (“*Delay 1.1*”), followed by a dotted line whose location indicated which of the two samples would be tested (“Cue”), followed by 4.5 sec of fixation (“*Delay 1.2*”), followed by a recognition probe (1 sec). Using a standard keyboard, participants made ‘Match’ (‘1’ key) /’non-match’ (‘2’ key) responses within a 2-sec window beginning with probe onset, with feedback displayed for remainder of response window (the fixation cross turned green for correct responses, red for incorrect responses). The ITI varied between 2-4 sec (**Fig. 1A)**. 50% of probes were matches of the cued item (e.g., the same face that had been shown during the sample period), and 50% of nonmatches were drawn from the same category as the cued item (e.g., a face that was different from the sample face). Throughout each block, the TMS coil was positioned to target area IPS2 in the right hemisphere, and randomization of memory set, cued category, and probe type were constrained so that spTMS was delivered on 50% of trials of each type. spTMS pulses were delivered 2-3 seconds after the offset of the cue, the unpredictable lag determined by selection (with replacement) from a flat distribution of lags that varied in 50 ms steps.

**Figure 1.**
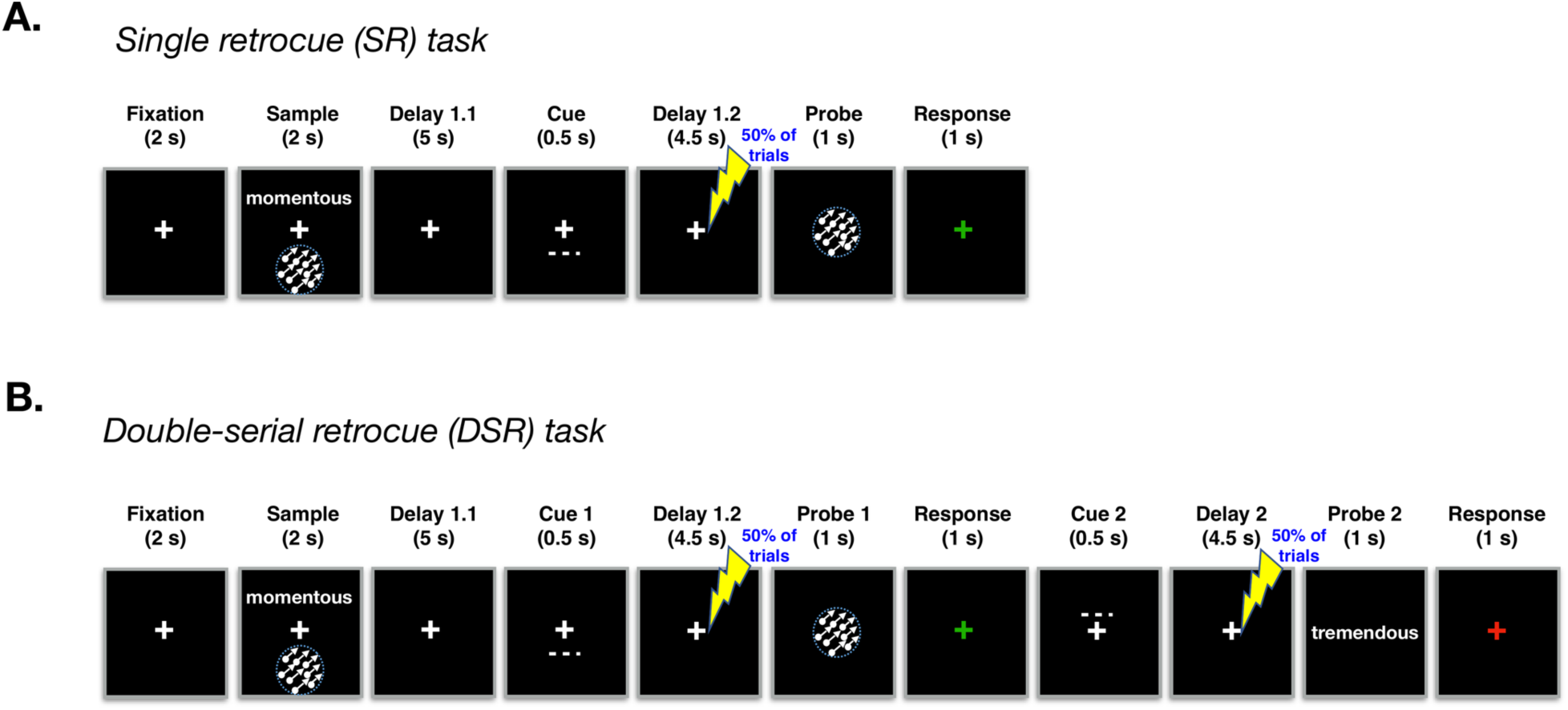
Behavioral task schematics. **A.** Schematic of the single-retrocue (SR) task. Participants memorized the identity of two sample items shown at separate locations and drawn from three possible categories (direction-of-motion, words, and faces (not shown)). Partway through the delay (i.e., at the end of *Delay 1.1*) a spatial retrocue indicated which of the two items would be tested, and during the ensuing *Delay 1.2* a single pulse of TMS (spTMS) was delivered on 50% of trials. At test, participants indicated whether the probe stimulus was a match or non-match of the cued memory item and received feedback. **B.** Schematic of the double-serial retrocue (DSR) task, the first half of which was identical to the SR task, followed by a second spatial retrocue (*Cue 2*) indicating which of the two samples would be tested by *Probe 2*, followed by a final delay during which spTMS was delivered with on 50% of the trials.

The double serial retrocue (DSR) task followed the procedure of the SR task except that the trial did not end with the first recognition probe. Instead, the 2-sec recognition window for “Probe 1” was followed by a second cue (“Cue 2”), followed by “*Delay 2*” (4.5 sec) and “Probe 2” (1 sec, plus additional 1 sec response capture window). Cue 2 appeared in the same location as Cue 1 on 50% of trials (“stay” trials), and spTMS was counterbalanced across all levels of Cue 1, Probe 1, Cue 2 (“stay” or “switch”) and Probe 2.

### Stimuli

The experimental stimuli were presented in MATLAB using the Psychophysics Toolbox (Brainard, 1997; Pelli, 1997; Kleiner et al., 2007) on an LCD with a resolution of 1920 x 1080 and background color set to black. The stimuli were viewed from a 70 cm viewing distance. The face stimuli were non-famous faces with neutral expressions selected from the set used by Rose et al. (2016; see also Rose et al., 2012). Matching face probes were exact matches. Nonmatching face probes were 50-70% morphs of the sample face and another face of the same gender. For word stimuli the critical feature was phonology: all probes were a different word from the sample but matches rhymed with the sample and nonmatches did not rhyme. Motion stimuli contained ∼125 white dots moving with 100% coherence at a speed of 3 deg/s for the duration of the stimulus presentation (2 sec), each trial in a direction chosen at random from the full 360-degree range. The direction of nonmatching motion probes differed from the sample within a range of 5-45 degrees.

### MRI data acquisition and preprocessing

Whole brain images were acquired with a 3T MRI scanner (Discovery MR750; GE Healthcare) at the Lane Neuroimaging Laboratory at the University of Wisconsin–Madison. High resolution T1-weighted images were acquired for all participants with an FSPGR sequence (8.2 ms repetition time (TR), 3.2 ms echo time (TE), 12° flip angle, 172 axial slices, 256×256 in-plane, 1.0 mm isotropic). The T1-weighted images were processed using the AFNI software program to align each participant’s brain with the MNI152_T1_1mm template. In AFNI, a mark was inserted in right intraparietal sulcus (rIPS2; coordinate: –22 70 58) and used as the target for spTMS (details below).

### spTMS targeting and delivery

rIPS2 was selected as a target for spTMS because of its role in implementing priority maps (e.g., Jerde et al., 2012), and in binding content to context in visual working memory (Gosseries, Yu, et al., 2018; Cai et al., 2019; Cai et al., 2020; Fulvio et al., 2023; Teng & Postle, 2024). It is important to note that because this study was not designed to test a hypothesis about a particular role for rIPS2 in the control of priority in working memory, our design did not require spTMS of a control site. Rather, rIPS2 was chosen as a target for spTMS for the theoretical reasons stated above, and because, in our experience, its stimulation was likely to produce a robust perturbation of working memory representations. The critical experimental controls in this study were the two epochs—*DSR Delay 2* and *SR Delay 1.2*—during which the spTMS-related effect on the uncued item was predicted to differ from its effects on the UMI (i.e., during *DSR Delay 1.2*).

Furthermore, as described in ***Behavioral tasks*** above, there was a 50% probability of spTMS delivery in each delay period, making spTMS delivery unpredictable on any given trial. As a result, both tasks included trials with no pulse and trials with one pulse; and additionally 25% of DSR task trials included two pulses, which critically mitigates concerns about differential arousal effects due to spTMS delivery between tasks and allows for a direct comparison of the spTMS effects between epochs to test our hypotheses. First, because participants could not anticipate how many pulses would occur on a given trial, any psychological arousal effect of an additional pulse in the DSR task would be an unpredictable, within-task occurrence rather than as a systematic difference between tasks. Second, we have shown in previous work that rTMS does not produce cumulative effects that might produce, e.g., a disparity in overall arousal (Hamidi et al., 2011).

spTMS targeting was achieved with a navigated brain stimulation (NBS) system that uses infrared-based frameless stereotaxy to coregister the location and position the participant’s head and that of the TMS coil according to the individual’s high-resolution MRI (NexStim eXimia, Helsinki, Finland). spTMS was delivered with an eXimia TMS Focal BiPulse transcranial magnetic stimulator fit with a figure-of-eight stimulating coil. NBS allows estimation of the electrical field induced by TMS at the targeted cortex using a model of the participant’s head and information about the coil position the distance from the coil to the cortical target. spTMS was delivered to the target to achieve an estimated intensity at the stimulation target of 90–110 V/m (60–75% of stimulator output, depending on the thickness of the participant’s scalp, cortex and depth of the target). The coil was oriented along the sagittal plane to induce an anterior-posterior direction of current, with individual adjustments to minimize EEG artifact. Stimulator intensity, coil position, and coil orientation were held constant for each participant for the duration of each session. To mask the sound of TMS coil discharge, participants were fitted with earbuds through which white noise was played during task blocks, with volume titrated such that the participants could not detect the click produced by coil discharge. Stimulation parameters were in accordance with published safety guidelines.

### EEG recording and preprocessing

EEG was recorded with a 60-channel cap and TMS-compatible amplifier, equipped with a sample-and-hold circuit that held amplifier output constant from 100 μs before stimulation to 2 ms after stimulation (NexStim eXimia, Helsinki, Finland). Electrode impedance was kept below 5 kΩ. The reference electrode was placed superior to the supraorbital ridge. Eye movements were recorded with two additional electrodes, one placed near the outer canthus of the right eye, and one underneath the right eye. The EEG was recorded between 0.1 and 350 Hz at a sampling rate of 1450 Hz with 16-bit resolution.

Data were processed offline using EEGLAB (Delorme & Makeig 2004) with the TMS-EEG signal analyzer (TESA) open-source EEGLAB extension (Rogasch et al., 2017; Mutanen et al., 2020) and Fieldtrip (Oostenveld, et al., 2011) toolboxes in MATLAB. We followed the TESA TMS-EEG analysis pipeline (http://nigelrogasch.github.io/TESA/). For delay periods for which no spTMS was delivered, a dummy spTMS event tag was added at a latency that matched the most recent spTMS-present trial delay period. Then, electrodes exhibiting excessive noise were removed and the data were epoched to –14 s to 4.5 s around the *Delay 1.2* spTMS event tag (for both the DSR and SR tasks) and –4.5 s to 4.5 s around the *Delay 2* spTMS event tag (for the DSR task only). The data were downsampled to 500 Hz. To minimize the TMS artifact in the EEG signal, the data were interpolated using a cubic function from –2 to 30 ms around the TMS pulse. This interpolation was also carried out on delay periods on which TMS was not delivered. Next, the data were bandpass filtered between 1 and 100 Hz with a notch filter centered at 60 Hz. Independent components analysis (ICA) was used to identify and remove components reflecting residual muscle activity, eye movements, blink-related activity, residual electrode artifacts, and residual TMS-related artifacts. Electrodes with excessive noise were interpolated using the spherical spline method. Finally, the data were re-referenced to the average of all electrodes that were included in the ICA.

### Decoding

A multivariate decoding analysis was used to address the primary question. The goal of the decoding analysis was to classify trials at the within-subject level according to whether or not an item was present in the memory array (at the category level, i.e., face, word, motion). Classification was carried out as a function of priority status during DSR *Delay 1.2*, during DSR *Delay 2*, and during SR *Delay 1.2*. Note that for timepoints prior to the cues, the items were coded according to the status that they would acquire. Thus, for each participant, 12 classifiers were trained to decode the presence or absence of each of 3 categories (face, word, motion), at each of 2 levels of priority (cued, uncued) and at each of 2 levels of spTMS (delivered (“present”), not delivered (“absent”)).

Classifiers were trained on spectrally transformed data. Spectral transforms of the preprocessed EEG data were calculated for each participant with the FieldTrip toolbox. A multi-taper-method convolution was performed on the epoched data using a fixed, Hanning-tapered window of 500 ms was used for integer frequencies from 2 to 50 Hz. This resulted in time-frequency representations consisting of 500 ms windows centered on timepoints spaced at 500 ms intervals, and epoched to the *Delay 1.2* spTMS pulse/dummy-code in both tasks, and to the *Delay 2* spTMS pulse/dummy-code in the DSR task, with the 12 data ‘present’ trial combinations transformed separately for each, and their 12 corresponding ‘absent’ trial combinations also transformed separately.

Before submitting the spectrally transformed data to classifier training, the power values were log10 transformed. The data were then temporally smoothed by averaging the power of each channel and frequency combination at each time point with the power at the preceding and following time point such that each time point aggregated 1.5 seconds of data. Because our interest was in the theta-, alpha-, and beta-frequency bands, we carried out a broadband classification including all three (4-30 Hz), and then we carried out a separate classification analysis in which the classifier was trained on individual frequency bands. Broadband classification was carried out at each timepoint. The feature set consisted of 60 channels x 3 frequencies (average theta power (4-8 Hz), average alpha power (8-13 Hz), and average beta power (13-30 Hz)). For the individual frequency band classification, we further divided the beta frequency band into low (13-20 Hz) and high (20-30 Hz), and each of the four frequency-band classifiers trained with a feature set of 60 channels. After frequency band averaging, the data were shuffled along the sample (trial) dimension and submitted to classification with approximately 120 samples provided per model. Classification used the MVPA-Light extension for the Fieldtrip toolbox (https://github.com/treder/MVPA-Light; Treder, 2020), using a logistic regression classifier with L2-regularization and lambda = 1. Five-fold classification was carried out with nested z-scoring per fold (i.e., the training data were z-scored and the resulting mean and standard deviation were used to z-score the test data to avoid leakage between the training and test data) and samples were averaged in random groups of two. In cases of uneven numbers of trials within classes, any remaining trials after averaging were discarded. In cases of imbalance across classes, the logistic model included a bias correction. Finally, the five-fold classification was repeated 10 times and the average area under the curve (AUC) was computed across all folds and repetitions for the particular model (combination of stimulus category, status, and TMS delivery). Here, AUC reflects how well the model can distinguish when a memory item was present versus absent from the EEG signal with higher AUC values signaling that the model is better at detecting when a participant was actively remembering something. These analyses were carried out using the HTCondor system deployed at the University of Wisconsin—Madison Center for High Throughput Computing, https://doi.org/10.21231/gnt1-hw21).

### Inferential statistical analyses

***Behavior.*** spTMS-related effects on behavior have previously been shown to occur for trials in which the uncued memory item was used as a lure but not for trials in which the probe was drawn from the cued category (Rose et al., 2016; Fulvio & Postle, 2020). Because the current study did not include lure probes, behavioral performance was not expected to be informative. Nevertheless, we analyzed recognition accuracy using two linear mixed effects models. The first analyzed behavioral performance for the first and second probe responses in the DSR task only with main effects of (1) probe response (two levels: first, second) (2) spTMS (delivered, not delivered); and (3) probe type (matching, non-matching) included in the analysis as fixed effects. Additionally, the interaction between probe response, spTMS, and probe type was included. All three fixed effects variables were coded as categorical variables. As random effects, the model included intercepts for the 12 participants. When we included participant-level random slopes for within-subject factors (e.g., spTMS), the model fits did not improve (e.g., with spTMS slope: ΔAIC = 3.9, *p* = 0.96), therefore, we did not use these more complex models. The second model included the same fixed effects but was fit to the data for all three probe responses (i.e., DSR task probes 1 & 2 and the single probe of the SR task). For this model, the probe response fixed effect had three levels (first (DSR), second (DSR), SR), and because the interaction was not significant, it was not included in the model. The *p*-values were obtained by *F*-tests for each term in the linear mixed-effects model.

***Decoding***. To analyze multivariate decoding effects of spTMS, we used cluster-based permutation tests on logistic regression performance as expressed as area under the ROC curve (AUC). The median AUC was computed for each participant, collapsed across the three stimulus categories, for each combination of priority status (PMI; UMI; IMI) and TMS delivery (spTMS-present; spTMS-absent). The data were then averaged at the group level across the 12 participants, and cluster-based permutation testing was used to identify timepoints of significant group-level classification performance against an AUC level = .5 (two-tailed) and, separately, to identify timepoints of significant differences in classification performance as a function of priority status. To address the primary question of this study, we report the results of these analyses within three task epochs during which spTMS could be delivered: DSR *Delay 1.2*, DSR *Delay 2*, and SR *Delay 1.2*. Additional follow-up between-task epoch comparisons were carried out on three timepoints: the spTMS delivery timepoint, and the first and second timepoints immediately following spTMS delivery. We report the results of these comparisons using one-tailed Wilcoxon signed-rank tests testing for greater classification AUC, at these time points, in DSR *Delay 1.2* than the other two task epochs. We also report the rank-biserial correlation coefficient (*rbc*) effect size.

### EEG dynamics

To gain insight about possible mechanisms underlying effects of spTMS on MVPA decoding, we also carried out analyses of phase and power dynamics in the EEG signal in relation to prioritization cues and to spTMS.

***EEG analyses of phase.*** All phase analyses relied on the results of a Hilbert transform. Prior to the transform, participants’ EEG signals were baseline-corrected with the 1000 ms preceding the trial (i.e., during the ITI) and downsampled to 100 Hz. Individual trial data were then averaged across channels. A Butterworth bandpass filter (order = 4) was applied to 4-second epochs centered on the prioritization cues and on spTMS pulses to isolate each of the four frequency bands that were used for decoding analyses: theta (4-8 Hz), alpha (8-13 Hz), low beta (13-20 Hz), and high beta (20-30 Hz). Filtering was implemented using Matlab’s filtfilt() function, which uses zero-phase forward and backward filtering. The filtered data were then analyzed using the Matlab function hilbert() and the instantaneous phase at each time point was computed from the analytic signal using the angle() function.

The resulting phase data were used to derive three measures of interest, each operating at a distinct level of analysis. The first, ***within-participant PLV***, quantifies phase consistency across trials within each individual, reflecting how reliably a participant’s neural phase aligned to a given event across repeated trials. The second, ***mean phase angle shift***, captures the direction of the event-induced phase change for each participant (averaged across trials), reflecting whether the event shifted phase in a consistent direction within individuals. The third, ***inter-subject PLV***, operates at the between-subject level and quantifies the consistency of those participant-level phase shifts across the group, indicating whether participants tended to show phase shifts in the same direction. Together, these three measures provide complementary perspectives on event-related phase reorganization: within-participant trial-to-trial consistency, the direction of individual-level phase changes, and the cross-participant alignment of those changes.

For prioritization cue-related impacts, we computed these measures relative to *Cue 1* of the DSR task and the single cue of the SR task. (We omitted data surrounding *Cue 2* of the DSR task from this analysis because a response and feedback period immediately preceded the cue, which would complicate the interpretation of the results.) For spTMS-related impacts on phase, we computed these measures for DSR *Delay 1.2*, DSR *Delay 2*, and SR *Delay 1.2*. We additionally computed these measures relative to the dummy-coded spTMS pulse in spTMS-absent trials. Because we could not know *a priori* when each frequency band might show a phase reset effect, we used a data-driven approach to identify an optimal analysis window for each combination of frequency band and event with a sliding time window selection approach. The window selection procedure utilized the combined set of all data around the two cues for the cue-related analysis and separately the combined set of all data around the three spTMS pulses for the spTMS-related analysis. The selected window was then used to compute each of the three measures for each condition separately. (Note that the window identified in spTMS-present trials was used in analyses of spTMS-absent trial data).

To identify the optimal post-event window via the sliding window approach, for cue-locked analyses, multiple candidate window lengths were defined, corresponding to integer numbers of oscillatory cycles (2–4 cycles for the theta and alpha bands, 3–6 cycles for the low and high beta bands). This ensured that lower frequencies were analyzed with longer windows and higher frequencies with shorter windows to preserve phase estimation reliability. For each candidate window, the “post-cue” period (0-500 ms relative to cue onset, cue duration = 500 ms) was sampled in 10 ms steps (corresponding to the 100 Hz sampling rate of the EEG data), and a pre-event window of equal length was defined immediately preceding the cue. The inter-trial phase-locking value (*PLV*) was computed for each participant in each post-cue window by averaging phase observations across all trials at each timepoint and then averaging all timepoints within the window. Separately, the corresponding baseline PLV was computed in the pre-cue window using the same procedure. Finally, pre-to-post differences were computed (*post-pre*). To select the optimal post-cue window for subsequent analyses, group-level *t*-statistics were computed for participants’ post – pre measures for each candidate window and cycle length. The window exhibiting the maximum absolute *t*-statistic was chosen as the representative post-cue window for that frequency band and used in the subsequent condition-specific analyses as described below. The window selection procedure for spTMS-related effects was identical, with the exception that the post-spTMS period began at 30 ms after the pulse to avoid the portion of data that was interpolated around the pulse during pre-processing (see above).

Having identified the analysis window for each frequency band and event-type (i.e., cue or spTMS) using the sliding-window, frequency-adaptive procedure, we computed the three measures within those windows for each participant and condition separately. For spTMS effects, event-related changes in the three measures were computed relative to a baseline corresponding to the same epoch from the spTMS-absent trial, such that observed effects reflected the impact of spTMS beyond spontaneous or task-related activity. Because our hypotheses related to between-condition differences, especially effects in low-beta, we primarily focused our inferential statistics on between-condition comparisons. For the within-participant event-induced PLV changes and group level mean phase shifts measures, we evaluated cross-condition differences (e.g., DSR *Cue 1* vs. SR *Cue*) within each frequency band using a paired permutation test with 5,000 iterations. In each iteration, the condition labels for each participant were randomly flipped to generate a null distribution of mean differences across participants. The two-tailed *p*-value was computed as twice the smaller proportion of permutations exceeding the observed difference. We also report the number of individual participants who demonstrated the group-level effect where these are interpretable. For the within-participant PLV, which is a scalar measure, sign-based individual counts are straightforward to interpret and we report the individual counts for significant contrasts. For the mean phase angle shifts, which is a circular measure, sign-based individual counts are only interpretable when the group-level mean difference does not approach the circular boundary (±180°) where the wrapping property of the circular distance computation means that trivially small individual differences in phase angle can flip the sign of the computed difference score, making sign-based counting unreliable as an index of individual consistency. This pattern illustrates precisely why group-level circular statistics are the appropriate inferential tool for these data because they are specifically designed to handle the geometry of circular distributions in a way that sign-based individual counts cannot.

To assess inter-subject PLV, reflecting the alignment of phase changes across participants rather than across trials, we computed the vector length of the mean phase shift across participants for each condition. This measure ranges from 0 to 1, where 0 indicates no consistency and 1 indicates perfect alignment of phase shifts across participants. Standard errors were estimated via 1,000 bootstrap resamples, and paired permutation tests (5,000 iterations) were used to test for cross-condition differences by randomly swapping condition labels across participants as was done for the other two measures. Finally, to account for multiple comparisons across frequency bands in these, *p*-values were FDR-corrected for the cue-related analyses and for the spTMS analyses.

Finally, we carried out time-resolved phase consistency analyses in 10 time windows from 50 ms to 500 ms post-cue/spTMS in 50 ms increments. At each time window, we computed each participant’s mean phase angle (averaged across trials) and tested whether these participant-level angles clustered significantly using the Rayleigh test, with the resultant vector length (*r*-value) quantifying clustering strength. Significant within-condition *r*-values indicating clustering of participant phases at the timepoints post-event were identified using the Rayleigh test with FDR correction across the 10 time windows. To test for between-condition differences for spTMS-present versus spTMS-absent trials, we used a permutation test with 1000 iterations of randomly swapping each participant’s condition labels (spTMS-present vs. spTMS-absent) and recomputing the *r*-value for each time point and condition and computed the difference between the *r-*values to form the null difference permutation distribution. The *p*-value was computed as twice the minimum of the proportions where the permuted difference was greater than or equal to, or less than or equal to, the observed difference (two-tailed test) and permutation *p*-values were FDR-corrected over the 10 time windows.

***EEG analysis of power.*** For each participant, for each channel and trial, we isolated the EEG signal for the 1 sec preceding the event of interest and, separately, for the 1 sec following the event. Individual trial data were then averaged across channels prior to analysis. Each channel-averaged trial was windowed using a Hanning taper to minimize spectral leakage. The power spectrum was computed using a fast Fourier transform (FFT) for each trial. From each trial’s resulting power spectrum, we extracted the relative power within the four frequency bands of interest as the proportion of the total spectral power (from 1-45 Hz) that fell within the target frequency band. For each participant, the mean power was computed across trials within each 1-second time window and each frequency band. Within-subject permutation tests (n = 5000) were used to assess whether observed pre vs. post changes were significant. These were carried out by randomly flipping ‘pre’ and ‘post’ labels across trials within each participant’s data set, maintaining equal proportions. Between-condition comparisons, in the case of comparing trials with spTMS delivery and those without, were also conducted using permutation tests (n = 5000) whereby all data were pooled and the label of ‘with’ and ‘without’ spTMS was randomly flipped, maintaining the original proportions.

## Results

### Recognition performance

The linear mixed effects model limited to DSR failed to find evidence that the delivery of spTMS had a significant effect on memory recognition accuracy (**Fig. 2**). In this analysis, only a significant probe x probe-type interaction was observed (*F*(1,88) = 4.046, *p* = .047), reflecting the fact that performance on probe 1 was similar for matching and non-matching probes (*M_matching_response1_* = .884 (*SD =* .049); *M_non-matching_response1_* = .893 (*SD =* .042), but performance on probe 2 was better for matching probes than for non-matching probes (*M_matching_response2_* = .871 (*SD =* .045); *M_non-matching_response2_* = .858 (*SD =* .054)). The second linear mixed effects model, which included SR task performance, also failed to find evidence of an effect of spTMS. In this analysis, only a main effect of probe was observed (*F*(2,139) = 5.717, *p* = .004), reflecting the fact that performance on the SR task and on Probe 1 of the DSR task (*M_SR_* = .883 (*SD =* .046); *M_DSR_response1_* = .889 (*SD =* .038)) were both better than performance on Probe 2 of the DSR task (*M_DSR_response2_* = .864 (*SD =* .045)).

**Figure 2.**
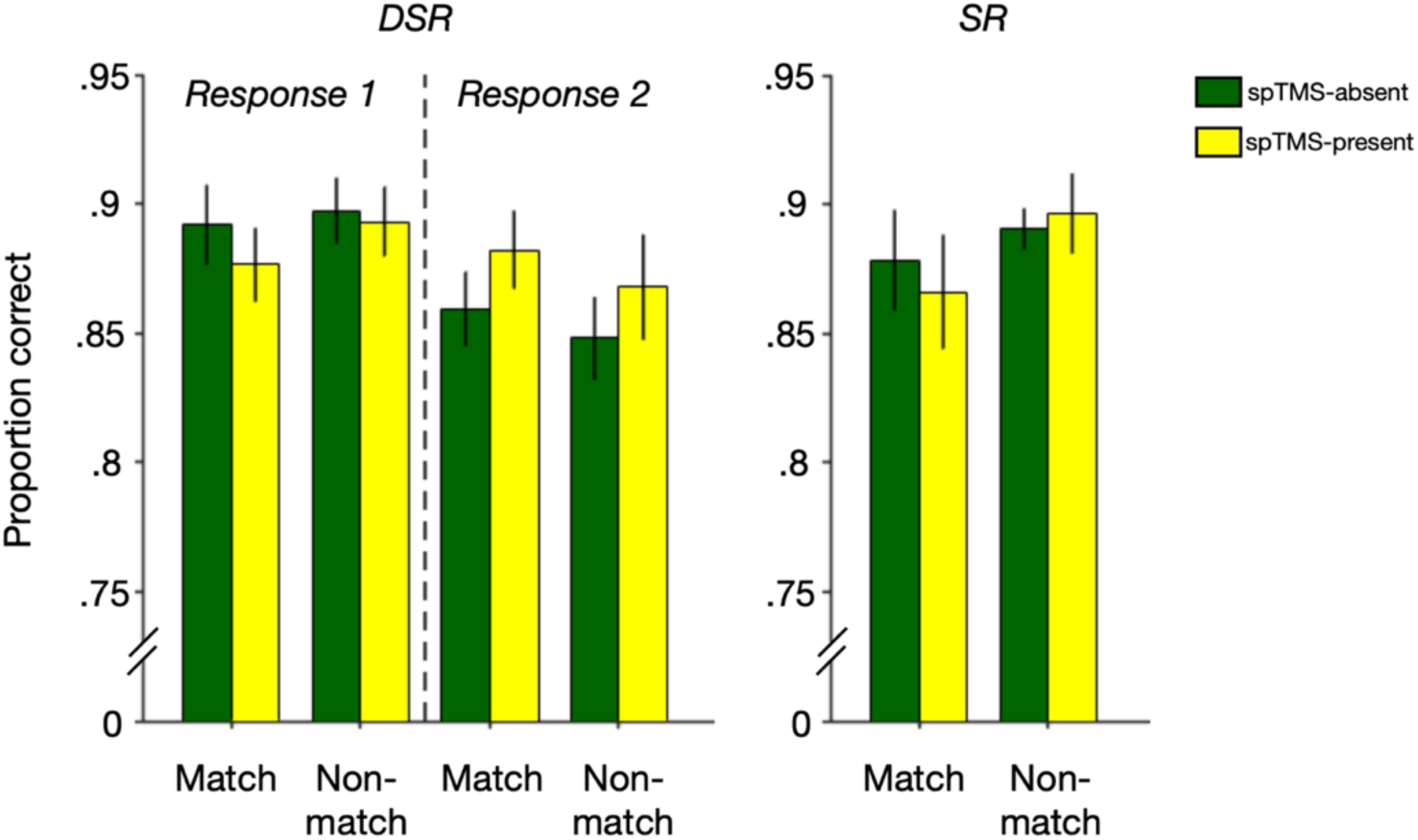
Recognition performance in the experimental tasks. Accuracy in the DSR and SR tasks as a function of probe type (match / non-match) and spTMS delivery. Error bars correspond to +/– 1 standard error of the mean for the n=12 participants.

### Decoding stimulus information from the broadband EEG signal

Beginning with *Delay 1.2* of the DSR task, prior to spTMS delivery neither item could be decoded above chance. For the UMI, classifier performance became significant at the time of spTMS delivery, and this effect persisted into the *Probe 1* epoch. For the PMI, classifier performance became significant one timepoint later than it had for the UMI, and this effect also persisted into the *Probe 1* epoch (**Fig. 3A**). For *Delay 2* of the DSR task, classification prior to spTMS was not different from chance for PMI or IMI, although decoder performance was significantly greater for PMI than for IMI. For the IMI, classifier performance rose above chance beginning with the second timepoint after spTMS (i.e., 1000 ms) and persisted into Probe 2; and for the PMI, classifier performance rose above chance at the time of spTMS and persisted into *Probe 2* (**Fig. 3B**). For the SR task, classifier performance did not differ from chance for either item during *Delay 1.2*. For the IMI, classifier performance rose above chance beginning with the second timepoint after spTMS and persisted into the Probe epoch; and for the PMI, classifier performance rose above chance one timepoint after spTMS and persisted into the Probe epoch (**Fig. 3C**). For trials in which spTMS was not delivered, UMI classifier performance never rose above chance, from the cue to response period, during either task and PMI classifier performance only rose above chance immediately before (*Delay 1.2*) or concurrent with (*Delay 2*) probe onset in the DSR task, and only briefly rose above chance in response to the retrocue in the SR task (**Fig. S1**). Thus, successful UMI/IMI decoding was observed only with spTMS delivery, and concurrent with spTMS only during *Delay 1.2* of the DSR task. One-tailed Wilcoxon signed-rank tests comparing UMI/IMI decoding at the spTMS timepoint confirmed significantly superior UMI classification performance for *Delay 1.2* relative to *Delay 2* (*W* = 63, *p* = .032, *rbc* = .615) and relative to SR *Delay 1.2* (*W* = 64, *p* = .026, *rbc* = .641; **Fig. 3D**). None of the comparable comparisons for subsequent timepoints (i.e., at a longer lag after spTMS) reached significance.

**Figure 3.**
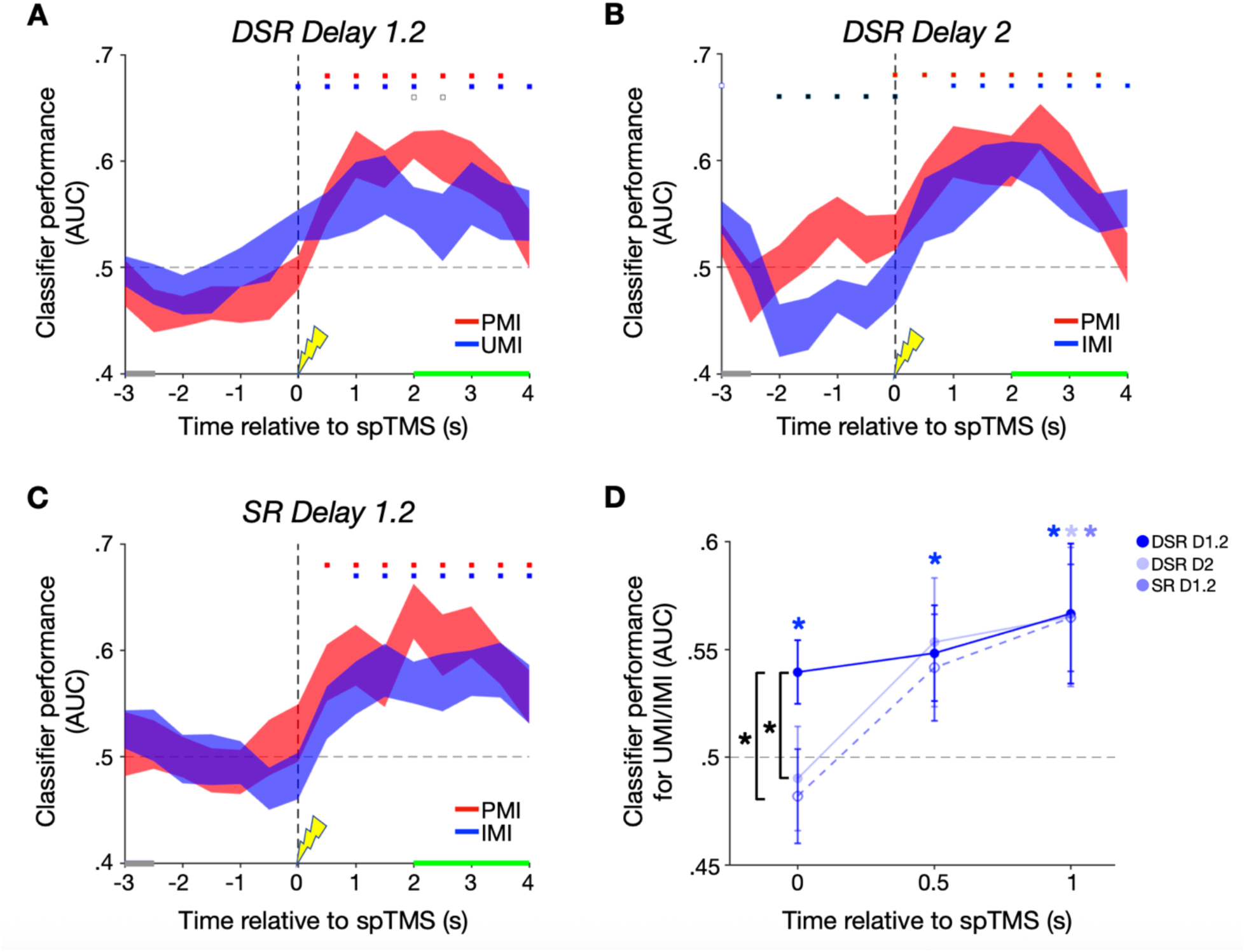
Broadband power classifier performance. **A.** Classifier AUC during *Delay 1.2* of the DSR task for the PMIUMI. Blue highlighting along the x-axis corresponds to the retrocue, green highlighting corresponds to the probe+feedback periods, and the spTMS pulse time-locked to 0 seconds. **B.** Same as **A.** for *Delay 2* of the DSR task. **C.** Same as **A.** & **B.** for *Delay 1.2* of the SR task. Error bands correspond to +/– 1 standard error of the mean for the n=12 participants. Red squares correspond to significant AUC clusters for PMI decoding; blue squares correspond to significant AUC clusters for UMI/IMI decoding; black squares correspond to significant AUC difference clusters between PMI and UMI/IMI. Filled markers: *p < .*05; empty markers: *p < .*1. **D.** Between-epoch comparison of UMI/IMI decoding for the three task epochs at the spTMS timepoint (0 s), the timepoint after spTMS delivery (.5 s), and two timepoints after spTMS delivery (1 s). Blue * markers: *p < .*05 based on cluster-based permutation test results in **A.**-**C.**; black * markers: *p* < .05 based on one-tailed Wilcoxon signed-rank tests. See **Figure S1** for classifier performance for trials without spTMS delivery.

*Decoding stimulus information from individual frequency bands.* Next, we decomposed the EEG data into classically defined frequency bands--theta ([4,8 Hz)), alpha ([8,13 Hz)), low beta ([13,20 Hz]), and high beta ([(20,30] Hz)—and repeated the analyses summarized in Figure 3. Inspection of the classification results across task epochs revealed no evidence for spTMS reactivation of the UMI/IMI in the theta band (**Fig. 4A**). The pattern of decoding from the alpha band matched what was seen in analyses of the broadband data: successful decoding of the UMI concurrent with the delivery of spTMS that was specific to *Delay 1.2* of the DSR task, with significant IMI decoding during *Delay 2* of DSR and *Delay 1.2* of SR beginning two time points later (**Fig. 4B**). One-tailed Wilcoxon signed-rank tests comparing UMI/IMI classification found significantly better classification performance for DSR *Delay 1.2* compared to DSR *Delay 2* (*W* = 73, *p* = .002, *rbc* = .872) at the spTMS timepoint, but no difference in classification performance when compared to SR *Delay 1.2* (*W* = 53, *p* = .151, *rbc* = .359; **Fig. 5A**). The overall pattern of decoding of the uncued item from the low beta band was similar the results from the alpha band, but with the emergence of significant decoding shifted one timepoint later for all trial types (**Fig. 4C**). However, one-tailed Wilcoxon signed-rank tests comparing UMI/IMI classification performance for DSR *Delay 1.2* with that of DSR *Delay 2* and SR *Delay 1.2* at each timepoint identified no significant differences (**Fig. 5B)**. Finally, decoding of the UMI/IMI from the high beta band was unsuccessful throughout DSR *Delay 1.2*, emerged late during DSR *Delay 2*, and emerged one timepoint after spTMS delivery in SR *Delay 1.2* (**Fig. 4D).** Despite the lack of statistically significant group-level decoding at the spTMS timepoint in the high beta band, one-tailed Wilcoxon signed-rank tests comparing UMI/IMI classification performance found significantly greater UMI classification performance for DSR *Delay 1.2* compared to IMI classification performance SR *Delay 1.2* at the spTMS timepoint (*W* = 65, *p* = .021, *rbc* = .667; **Fig. 5A**). Comparisons of the DSR *Delay 1.2* classification performance with the other epochs at the two timepoints post-spTMS did not identify any differences. However, a post-hoc comparison found significantly greater IMI classification performance for DSR *Delay 2* compared to SR *Delay 1.2* (*W* = 61, *p* = .046, *rbc* = .564). As with the broadband analysis, no successful frequency band-specific decoding of the UMI or IMI was obtained on trials without spTMS delivery (**Figure S2**).

**Figure 4.**
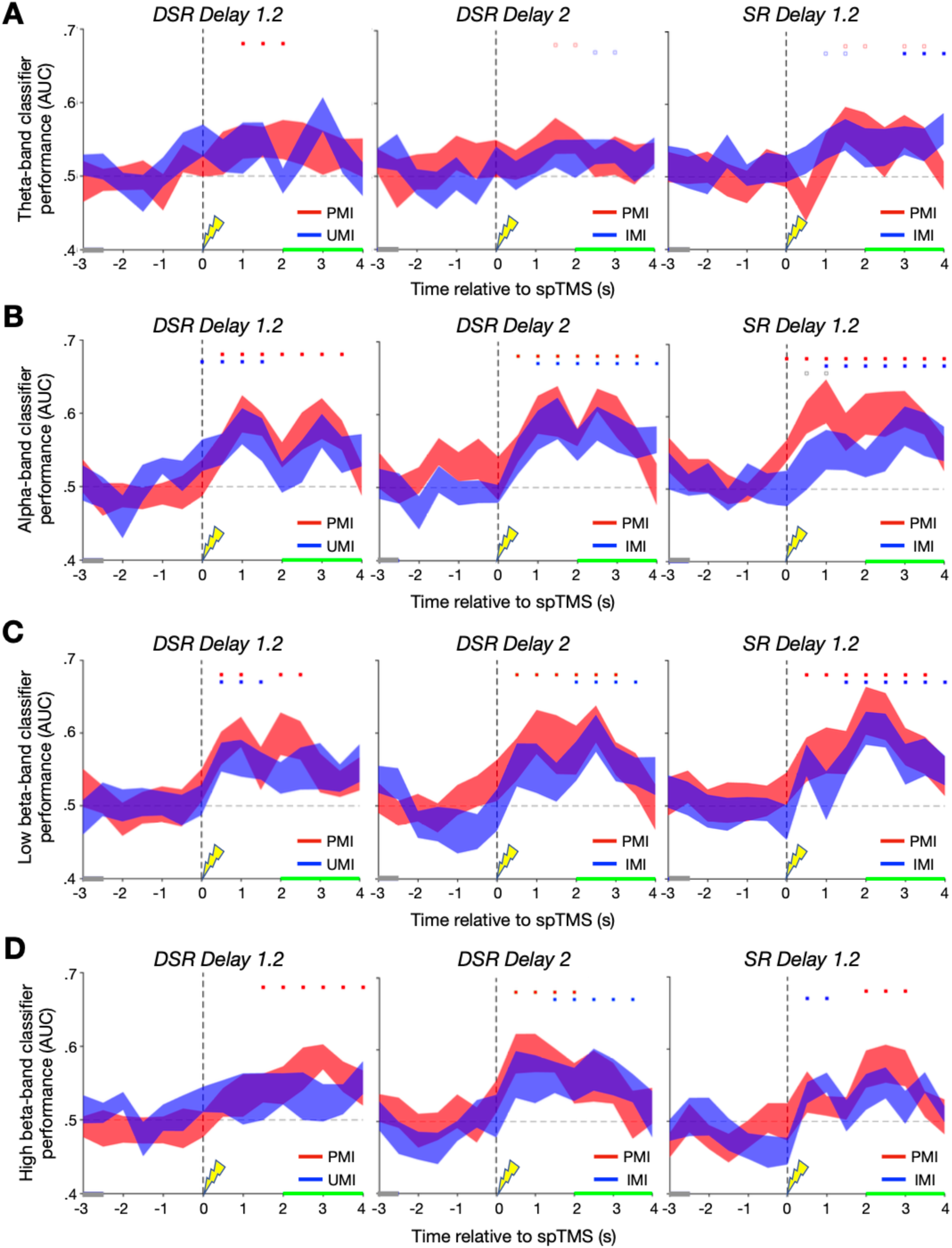
Frequency-band specific classifier performance. **A.** Theta-band Classifier AUC during the DSR *Delay 1.2* epoch (left), DSR *Delay 2* (middle), and SR *Delay 1.2* (right) for the prioritized memory category (PMI) in red and the unprioritized/irrelevant memory category (UMI/IMI) in blue. Gray highlighting along the x-axis corresponds to the retrocue period, green highlighting corresponds to the probe+feedback period, with the delay period occurring between and the spTMS pulse time-locked to 0 seconds. **B.** Same as **A.** for alpha-band classifier AUC. **C.** Same as **A.** & **B.** for low beta-band classifier AUC. **D.** Same as **A.**, **B.**, & **C.** for high beta-band classifier AUC. Error bands correspond to +/– 1 standard error of the mean for the n=12 participants. Red significance squares correspond to significant AUC clusters for PMI decoding; blue squares correspond to significant AUC clusters for UMI/IMI decoding; black squares correspond to significant AUC PMI-UMI/IMI difference clusters. Filled markers: *p < .*05; empty markers: *p < .*1. See **Figure S2** for classifier performance for trials without spTMS delivery.

**Figure 5.**
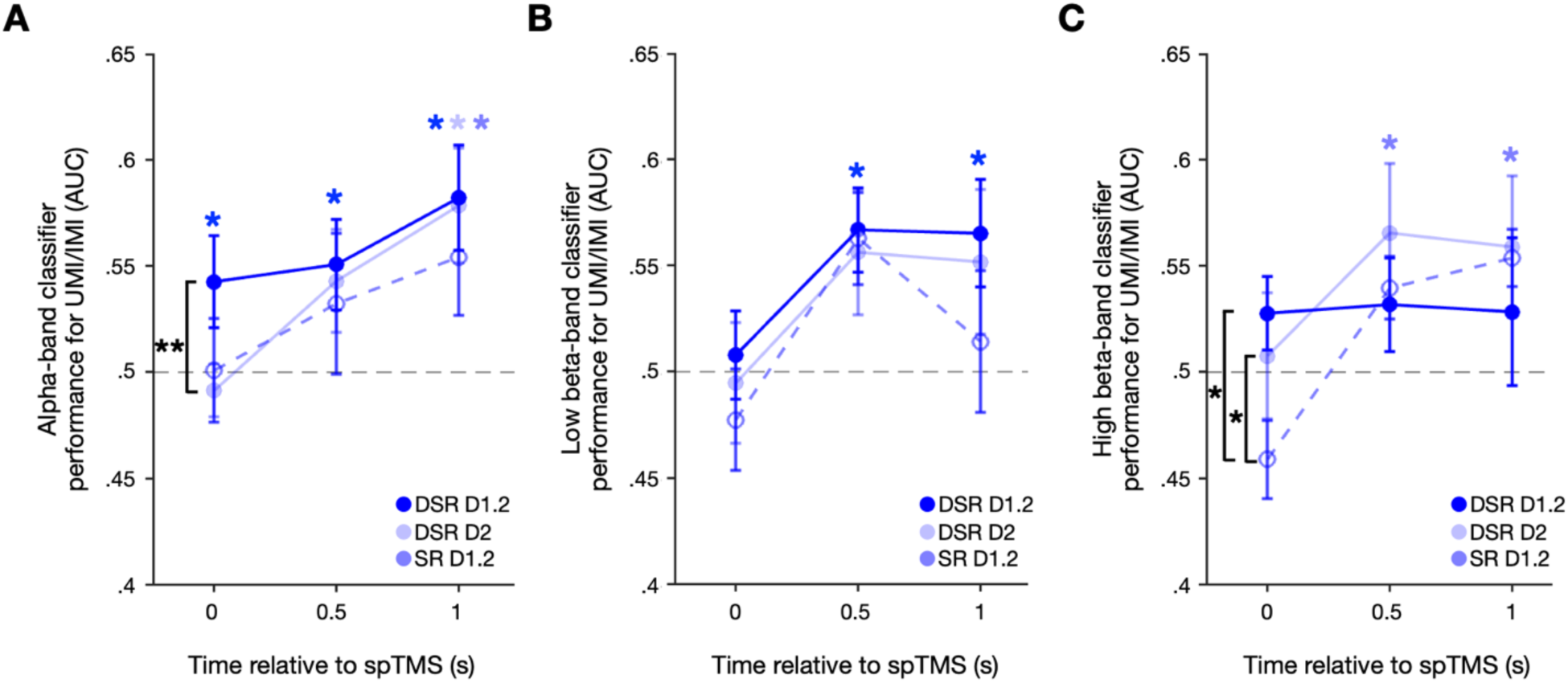
UMI/IMI classification performance comparisons. **A.** Between-epoch comparison of alpha-band UMI/IMI decoding for the three task epochs at the spTMS timepoint (0 s), the timepoint after spTMS delivery (.5 s), and two timepoints after spTMS delivery (1 s). **B.** Same as **A.** for low beta-band UMI/IMI decoding. **C.** Same as **A.** and **B.** for high beta-band UMI/IMI decoding. Blue * markers: *p < .*05 based on cluster-based permutation test results in **A.**-**C.**; black * markers: *p* < .05; black ** markers: *p* < .01 based on one-tailed Wilcoxon signed-rank tests.

### Interim Discussion

Whereas there was no evidence for above-chance decoding of either the PMI or the UMI during the early portion of any delay period, patterns of decoding in the second half of delay periods varied with priority status (i.e., UMI vs. IMI vs. PMI). For the uncued item, successful decoding of the UMI returned concurrent with the delivery of spTMS (i.e., during DSR *Delay 1.2*), whereas for the IMI successful decoding lagged by at least two timepoints during (i.e., during DSR *Delay 2* and SR *Delay 1.2*). When the data were decomposed into classical frequency bands, this priority-specific effect was seen in the alpha band and, with a one-time-window lag, in the low beta band. For the cued item (the PMI) the return of above-chance decoding during *Delay 1.2* of the DSR task lagged the UMI by one time window in the analyses of broadband and alpha band-filtered data. Although it is possible that these results reflect specific “reactivation” effects of spTMS on the UMI, this interpretation is complicated by two factors. First, pre-spTMS decoding of the PMI was at chance, and generally not different from the UMI, thus failing to demonstrate that the two items were, indeed, held at different levels of priority. Second, the fact that decodability of both items tended to ramp up late in the delay period in many conditions might be consistent with a nonspecific strengthening of stimulus information in anticipation of the recognition probe. For these reasons, it was important to assess whether the retrocues and spTMS had UMI-specific effects on aspects of the EEG data that may explain the patterns of MVPA decoding—specifically phase and power dynamics of the EEG signal. Evidence that these measures also responded differentially for the UMI would support the interpretability of the differential decoding effects and might also provide insights into the neurophysiological bases of the “spTMS reactivation” effect.

### EEG dynamics

#### Phase dynamics

*Prioritization cues.* The cue presented during DSR *Delay 1* (which designated the UMI) was associated with a significant increase in within-participant PLV in the low-beta band when compared with the effects of the cue from the SR task (which designated the IMI; *p* = .044, FDR-corrected; **Fig. 6A**). Consistent with the group-level effect, 10/12 participants showed greater low-beta within-participant PLV for DSR Cue 1 than the SR cue. Accompanying this effect, mean phase angle shifts differed significantly between *DSR Delay 1* and *SR Delay 1* in the alpha, low-beta, and high-beta bands (all *p*-values < .001, FDR-corrected; **Fig. 6B**): in theta, the difference was the result of a larger shift in response to the cue during SR *Delay 1*; in alpha and low-beta, the differences were the result of phase shifts in opposing directions in DSR *Delay 1* vs. SR *Delay 1*. Of the significant phase shift contrasts, only the cue-related low-beta effect met the criterion for interpretable counts (see Methods – ***EEG analyses of phase*** for more information): 8/12 participants’ data were in the group-level direction with a group mean difference = 136°. By contrast, the cue-related alpha effect showed near-chance consistency across participants (alpha: 6/12, group mean difference = 118°) and the cue-related high beta effect fell near the ±180° circular boundary (group mean difference = 179°).

**Figure 6.**
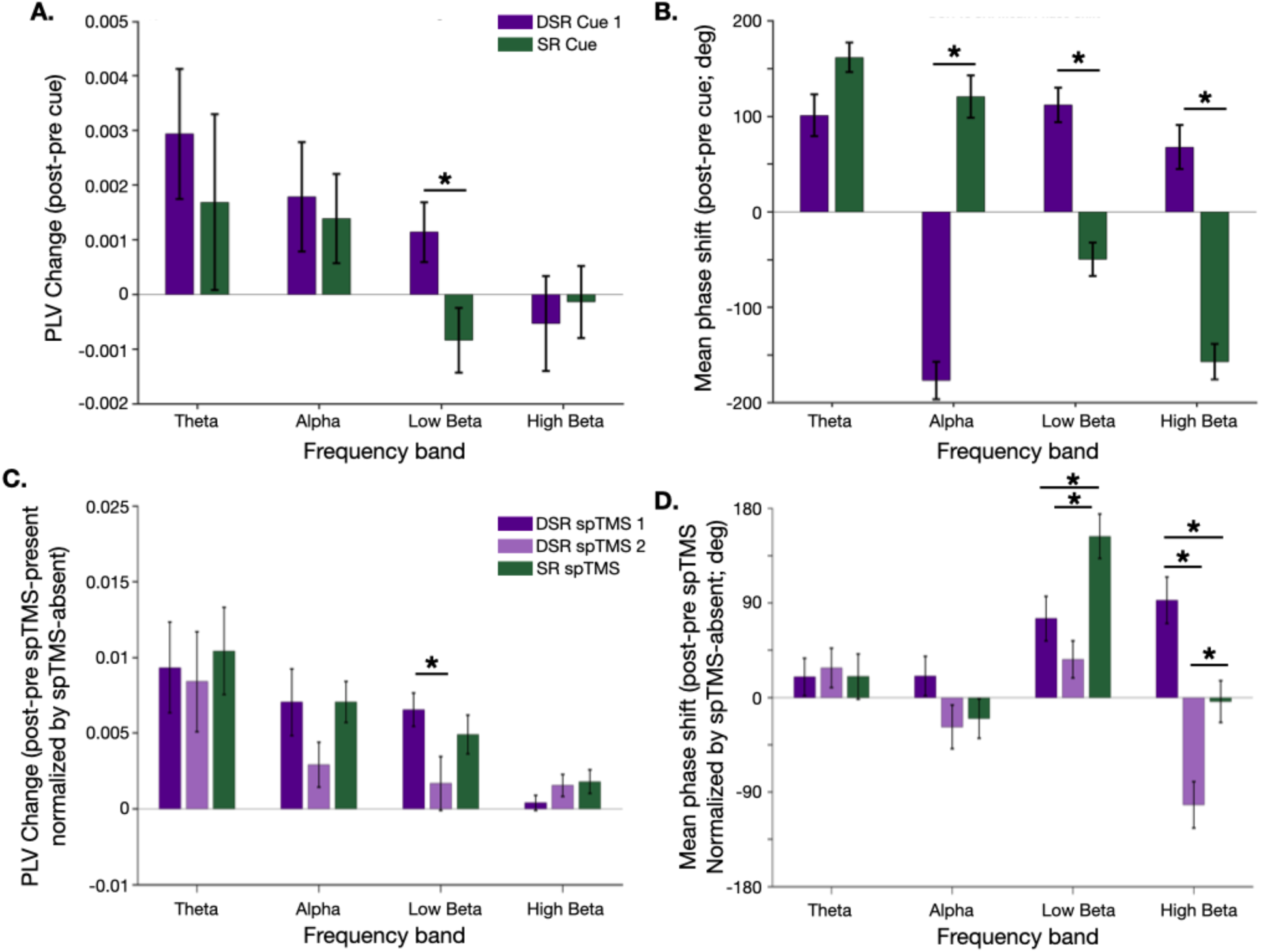
Analysis of phase dynamics. **A.** Cue-evoked within-participant PLV (post – pre PLV) for each of four frequency bands in response to *Cue 1* during DSR *Delay 1* (indigo) and the cue during SR *Delay 1* (green). **B.** Cue-evoked mean phase angle shift for each of four frequency bands, in response to *Cue 1* during DSR *Delay 1* (indigo) and the cue during SR *Delay 1* (green). **C.** Same as **A.,** but for spTMS-evoked within-participant PLV, normalized by spTMS-absent trials, for DSR *Delay 1.2* (indigo), DSR *Delay 2* (transparent indigo), and SR *Delay 1.2* (green). **D.** Same as **B.**, but for spTMS-evoked mean phase angle shift (normalized) in DSR *Delay 1.2* (indigo), DSR *Delay 2* (transparent indigo), and SR *Delay 1.2* (green). * markers: *p* < .05, FDR-corrected; error bars correspond to +/− 1 *SEM.* See **Figures 6-1 & 6-2** for the post-cue and post-spTMS time-resolved phase consistency analyses, respectively and **Figure 6-3** for cue– and spTMS-evoked changes in oscillatory power.

No other significant differences in cue-related effects across tasks and frequency bands were observed for within-participant PLV, mean phase angle shift, or inter-subject PLV (all *p*-values ≥ .1853, FDR-corrected). Furthermore, time-resolved analyses showed no evidence of between-participant phase clustering at any post-cue timepoint (**Fig. 6-1**).

*spTMS.* In general, spTMS-related effects on within-participant PLV were markedly larger in magnitude in comparison to cue-related effects. The only difference in these effects across conditions was that in the low-beta band. spTMS triggered larger within-participant PLV during DSR *Delay 1.2* than during DSR *Delay 2* (*p* = .026, FDR-corrected; for all other contrasts *p*-values ≥ .059, FDR-corrected; **Fig. 6C**). Consistent with this effect, 10/12 participants showed greater normalized PLV during DSR Delay 1.2 than DSR *Delay 2*, and 9/12 showed greater normalized PLV during SR *Delay 1.2* than DSR *Delay 2*. The non-significant contrast between DSR *Delay 1.2* and SR *Delay 1.2* showed near-chance consistency among participants (6/12), as expected.

For group-level spTMS-evoked mean phase angle shifts, significant differences between conditions were found in the low-beta and high-beta bands (**Fig. 6D**). In low-beta, the effect was significantly larger in the SR *Delay 1.*2 than in DSR *Delay 1.2* and DSR *Delay 2* (both *p*-values ≤ .0198, FDR-corrected), whereas in high-beta, it was significantly larger during DSR *Delay 1.2* than during DSR *Delay 2* and SR *Delay 1.2* (both *p*-values ≤ .0048, FDR-corrected) and significantly larger during DSR *Delay 2* than during SR *Delay 1.2* (*p* = .0048, FDR-corrected). Only the spTMS-evoked high-beta contrast between DSR *Delay 1.2* and DSR *Delay 2* met the criterion for interpretable counts (10/12; group mean difference = 74°).

Inter-subject PLV changes were not significantly different across any conditions within any frequency bands (all *p*-values ≥ .4572, FDR-corrected; results not depicted). However, time-resolved analyses revealed post-spTMS-specific between-participant phase clustering in all conditions that was absent in spTMS-absent trials. Clustering emerged in all four frequency bands following spTMS, but with distinct temporal profiles: the clustering was persistent throughout the analysis time range (i.e., 500 ms post spTMS) in theta and alpha, but transient in low-beta and high-beta. (**Fig. 6-2)**.

To summarize, at the level of individual phase dynamics, the effect of the cue and of spTMS were specific to the low-beta band. At the group level, the cue-related mean phase angle shift for DSR *Cue 1* vs. SR *Cue* was opposite in sign in the low-beta beta band in comparison to the theta and alpha bands, and a complex pattern of spTMS-related mean phase angle shifts was observed in the low– and high-beta bands.

#### Power dynamics

To address the possibility that the observed phase effects were driven by amplitude changes rather than genuine phase reorganization, we tested for condition-specific increases in oscillatory power evoked by the cues and/or spTMS. It was uniformly the case that power declined over time, whether assessed in response to the cues (**Fig. 6-3**), or in the post-cue delay periods on spTMS-present and spTMS-absent trials (**Fig. 6-3**). The magnitude of these declines generally decreased with increasing frequency. These dynamics are characteristic of active maintenance during working memory delays and indicate that the phase-related effects reported above were not confounded by systematic changes in oscillatory power.

## Discussion

The results presented here demonstrate patterns of spTMS-related decoding and of EEG dynamics that provide insight into the representation of priority in working memory. For the uncued item, only during DSR *Delay 1.2*, when it had the status of UMI, was the recovery of decoding concurrent with spTMS, and thus plausibly linked to the instantaneous effects of the perturbation. In the other two delay periods, recovery of decoding of the IMI occurred several timesteps later, whereas for the cued item (the PMI) the latency of spTMS-related decoding was comparable across conditions. Frequency band-specific analyses revealed the alpha and low-beta bands to support decoding of all items in all instances where broadband decoding had been observed. Analyses of phase dynamics in the EEG indicated that only during DSR *Delay 1.2* did the prioritization cue and spTMS produce phase shifts in the low-beta band at the within-participant level.

These results join a large body of evidence indicating that the deprioritization of information is accomplished via a representational transformation that distinguishes it from prioritized information, as well as from information that is no longer needed in working memory (e.g., Lewis-Peacock et al., 2012, van Loon, 2018, Yu, Teng, & Postle, 2020; Wan, 2020; Fulvio & Postle, 2020; Wan et al. 2024). They extend this literature by bringing into focus the idea that beta-band dynamics may play an important role in the encoding of priority. The first demonstration of spTMS-triggered decoding of the UMI isolated this effect to the beta band (Rose et al., 2016). More recently, it has been shown that when the EEG signal on the scalp is decomposed into a set of discrete coupled oscillators, it is components in the beta band whose sensitivity to spTMS correlates with spTMS-related declines in behavior. Here, we provide evidence for a specific role for phase dynamics in the low-beta band at the within-participant level, in that only the cue designating the UMI is associated with a phase shift in low-beta, and the effect of spTMS that is unique to DSR *Delay 1.2* is a phase shift in low-beta.

The delivery of single pulses of transcranial magnetic stimulation (spTMS) to posterior parietal cortex while the EEG is simultaneously recorded has revealed the “natural frequency” of this region to be in the low-beta range (Rosanova et al., 2009, Kundu et al., 2014), in comparison to alpha when spTMS targets occipital cortex, and high-beta/low-gamma when spTMS targets premotor cortex (Rosanova et al., 2009) and PFC (Ferrarelli et al., 2012). Furthermore, whereas spTMS-evoked phosphene perception is predicted by pre-TMS power and phase in spontaneous alpha when spTMS targets occipital cortex, it depends on pre-TMS power in spontaneous low-beta when PPC is targeted (Samaha et al., 2017), and global mean field amplitude triggered by spTMS of PPC fluctuates with instantaneous phase of the beta oscillation (Kundu et al., 2015). In the present study, and in Fulvio et al. (2024), spTMS was delivered to IPS2. Building from this, our tentative proposal is that priority in working memory may be encoded in a frontoparietal priority map that harnesses endogenous oscillations in the low-beta band to maintain the uncued item in an unprioritized state. (Note that, for reasons summarized below, the anatomical element of this idea will require future research designed to test hypotheses about anatomical specificity.) From this perspective, the fact that spTMS produces involuntary retrieval of the UMI at the behavioral level (Rose et al., 2016; Fulvio & Postle, 2020) suggests that the spTMS-triggered changes in low-beta band dynamics that are specific to the UMI may reflect a disruption of its low beta-encoded priority status. Importantly, our current construal of this mechanism is that it is different from the proposed role for beta bursts in the implementation of top-down inhibitory control (e.g., Wessel & Anderson, 2024; Lundqvist et al., 2024), for two reasons. First, decoding of the UMI from the beta band is sustained for periods of time much longer than what would be expected from a phasic signal carried by a burst. Second, the delivery of spTMS in these studies is unpredictable, and so unlikely to be tightly synchronized with an endogenously formulated, and brief, control signal.

There are many experimental applications for TMS in working memory research, including the disruption or enhancement of function, testing hypotheses about the contribution of one or more brain regions to a mechanism of interest, and testing hypotheses about task-related connectivity, and these often require the inclusion of an active stimulation control site to support strong inference (Johnson, et al., 2021). For the present study, however, because our empirical hypotheses related to the oscillatory bases of the “spTMS reactivation effect” (Rose et al., 2016, Fulvio & Postle 2020), and our theoretical interest to mechanisms of (de)prioritization, our design did not require an anatomically distinct control site for spTMS. Rather, the critical controls were spTMS-absent epochs of DSR *Delay 1.2*, plus spTMS-present and –absent conditions for epochs during which the uncued item was not deprioritized (i.e., DSR *Delay 2* and SR *Delay 1.2*). Thus, although our design used spTMS, there is a sense in which it can be conceptualized as using a logic similar to experiments presenting a photic “ping” stimulus during the delay period, to test a hypothesis about, for example, stimulus decodability or representational format (c.f., Wolff et al., 2015; Kandemir et al., 2025).

One concern that might apply more so to spTMS than to other experimental interventions, such as photic “pinging,” is uneven influences on physiological arousal. However, the fact that oscillatory power showed a general decrease following both cue and spTMS events, across all frequency bands and across all conditions, rules out the concerns about possible condition-specific arousal responses. This also rules out the possibility that the true basis of the condition-specific effects in phase dynamics that we report here may have been changes in oscillatory power.

