## Supplementary material for "Effects of TMS on the decoding and electrophysiology of priority in working memory": Figure 3-1

**Supplementary Data**

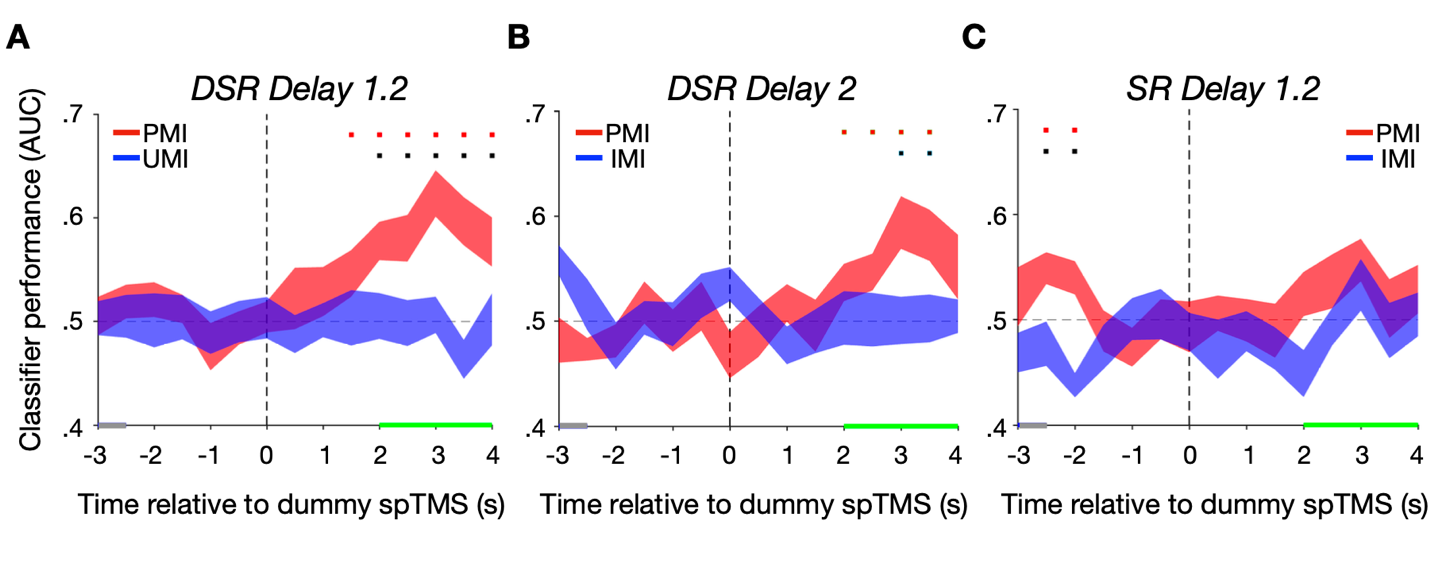

**Figure 3-1. Broadband power classifier performance for trials without spTMS. A.** Classifier AUC during the DSR Delay 1.2 epoch for the prioritized memory category (PMI) in red and the unprioritized/irrelevant memory category (UMI/IMI) in blue. Gray highlighting along the x-axis corresponds to the retrocue period, green highlighting corresponds to the probe+feedback period, with the delay period occurring between and the spTMS pulse time-locked to 0 seconds. **B.** Same as **A.** for DSR Delay 2. **C.** Same as **A.** & **B.** for SR Delay 1.2. Error bands correspond to +/– 1 standard error of the mean for the n=12 participants. Red significance squares correspond to significant AUC clusters for PMI decoding; blue squares correspond to significant AUC clusters for UMI/IMI decoding; black squares correspond to significant AUC PMI-UMI/IMI difference clusters. Filled markers: *p < .*05; empty markers: *p < .*1. Blue * markers: *p < .*05 based on cluster-based permutation test results in **A.**-**C.**

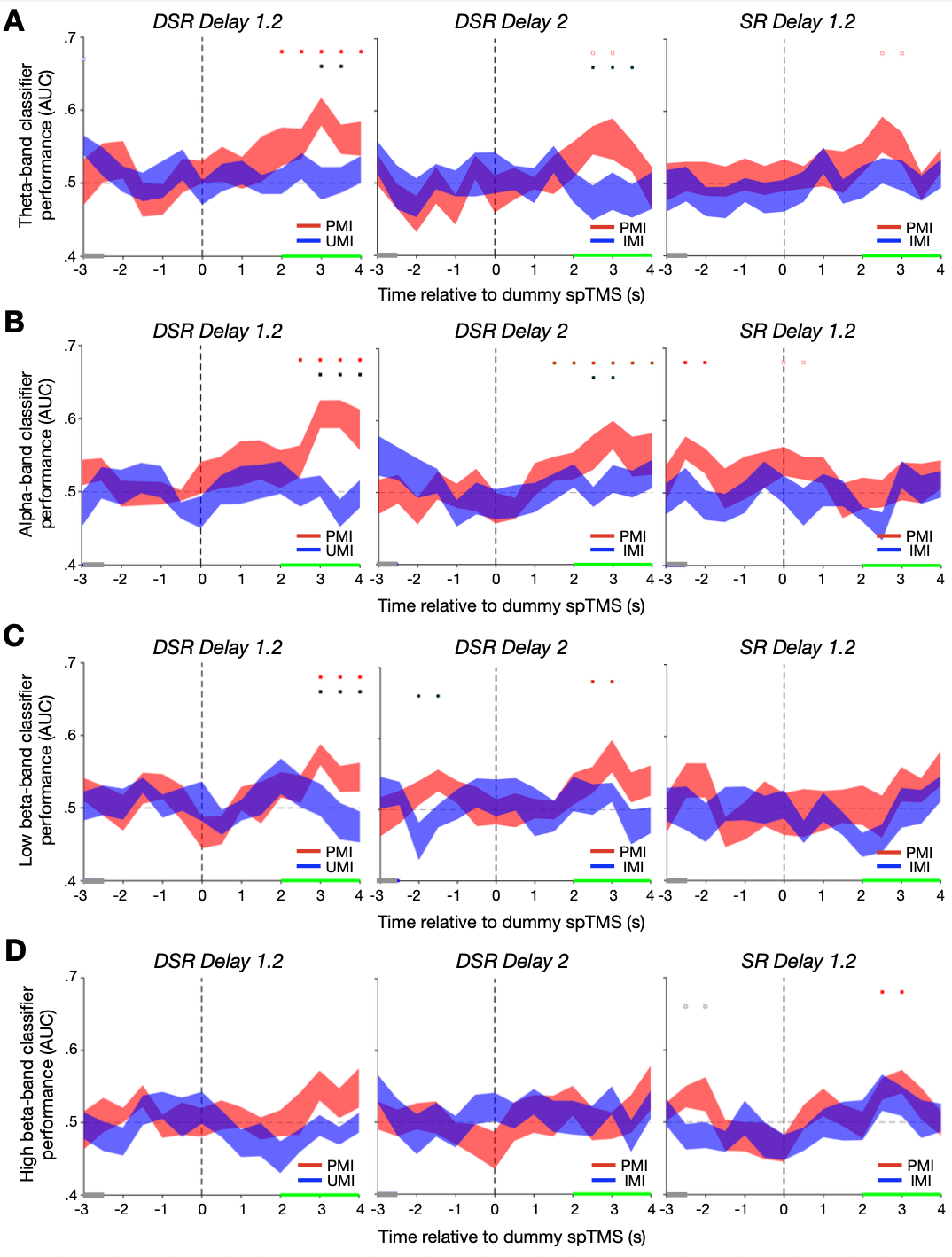

**Figure 4-1. Frequency-band specific classifier performance for trials without spTMS delivery. A.** Theta-band Classifier AUC during the DSR Delay 1.2 epoch (left), DSR Delay 2 (middle), and SR Delay 1.2 (right) for the prioritized memory category (PMI) in red and the unprioritized/irrelevant memory category (UMI/IMI) in blue. Gray highlighting along the x-axis corresponds to the retrocue period, green highlighting corresponds to the probe+feedback period, with the delay period occurring between and the spTMS pulse time-locked to 0 seconds. **B.** Same as **A.** for alpha-band classifier AUC. **C.** Same as **A.** & **B.** for low beta-band classifier AUC. **D.** Same as **A.**, **B.**, & **C.** for high beta-band classifier AUC. Error bands correspond to +/– 1 standard error of the mean for the n=12 participants. Red significance squares correspond to significant AUC clusters for PMI decoding; blue squares correspond to significant AUC clusters for UMI/IMI decoding; black squares correspond to significant AUC PMI-UMI/IMI difference clusters. Filled markers: *p < .*05; empty markers: *p < .*1.

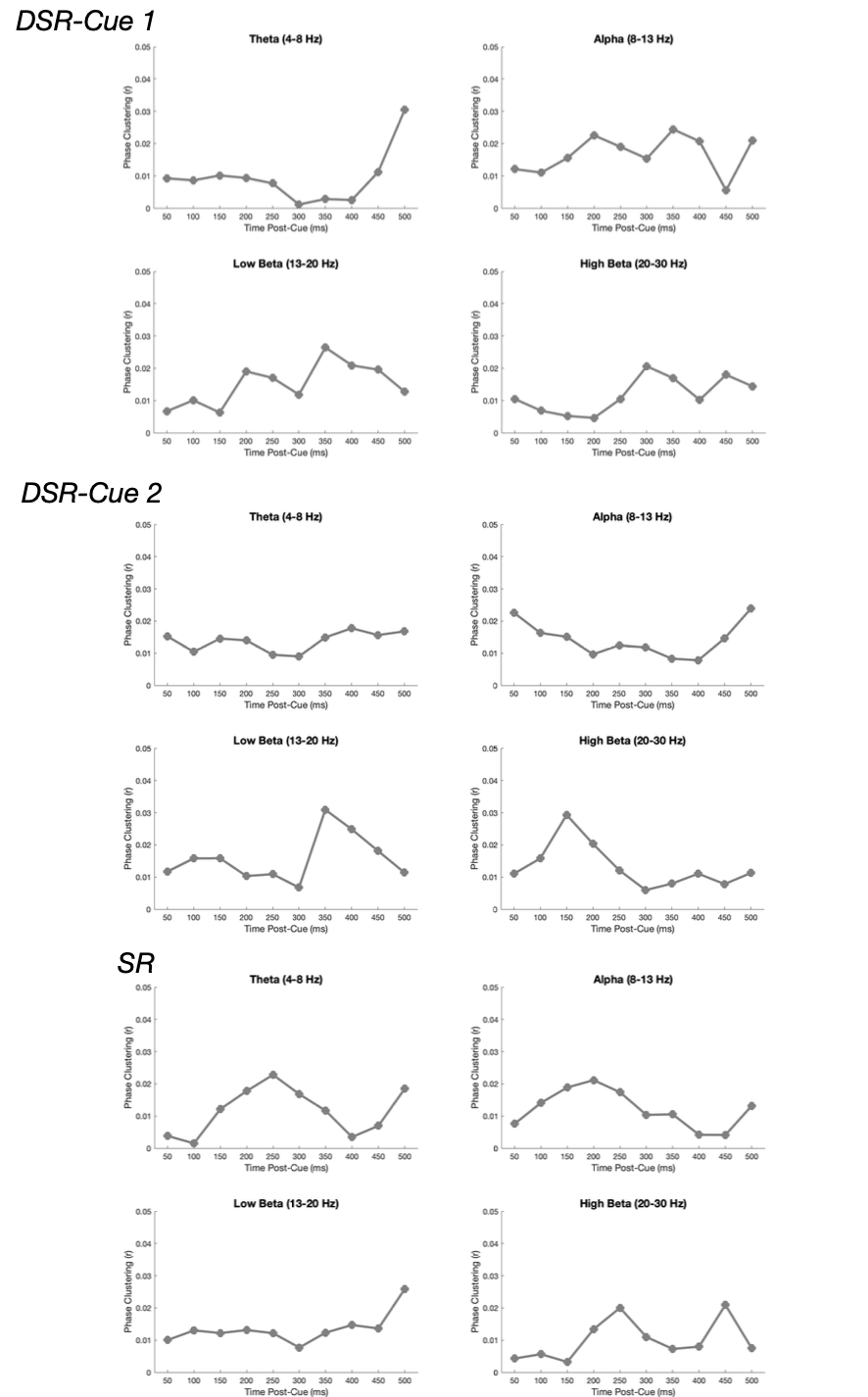

**Figure 6-1. Time-resolved post-cue phase consistency analysis.** Post-cue phase consistency between participants at 10 timepoints 50 ms apart for each of the four frequency bands and three retrocues. None of the consistencies reached significance after FDR correction.

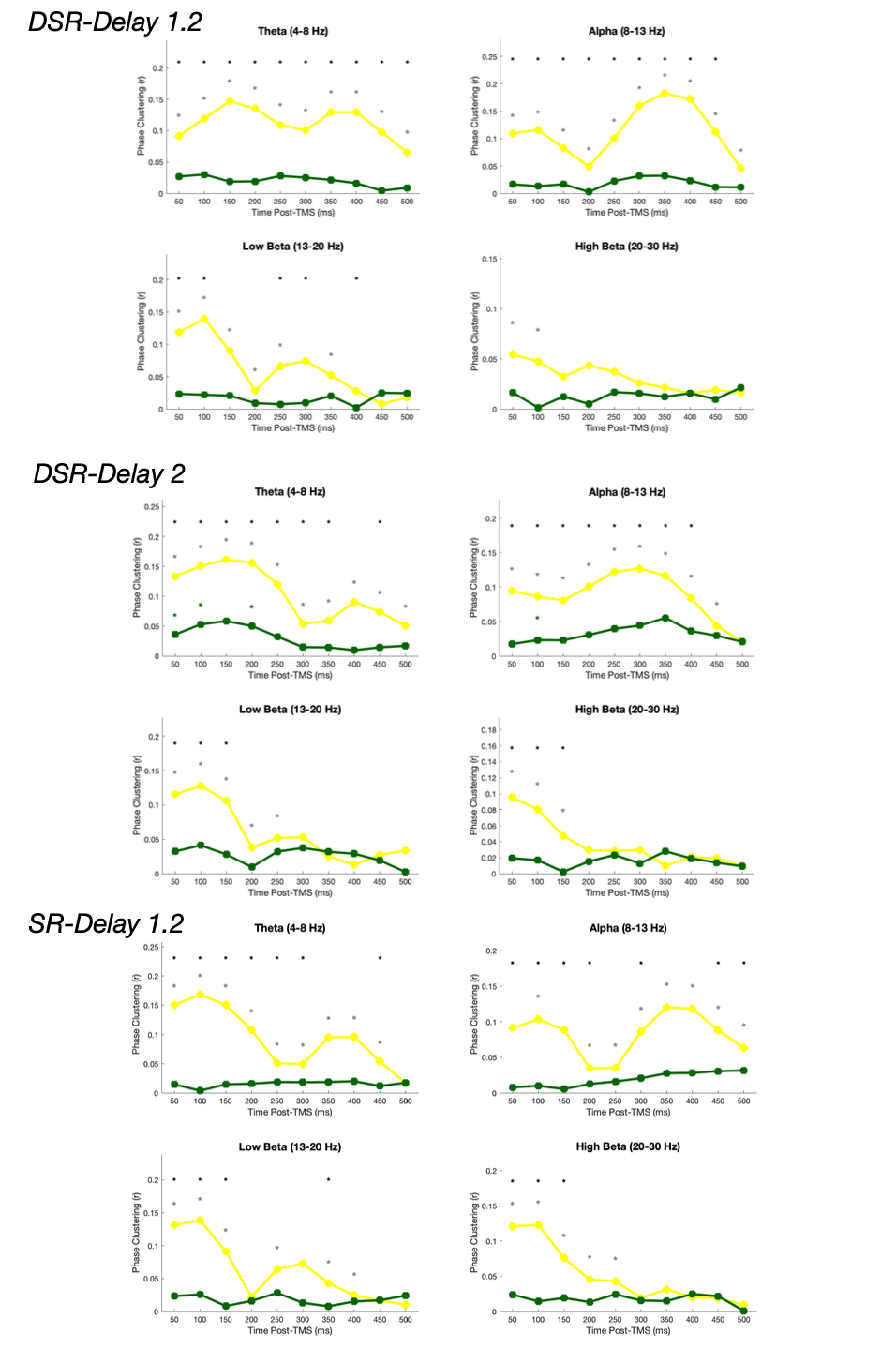

**Figure 6-2. Time-resolved post-spTMS phase consistency analysis.** Post-spTMS phase consistency between participants (yellow) at 10 timepoints 50 ms apart for each of the four frequency bands and three retrocues. For comparison, post-dummy spTMS phase consistency is plotted in green. Significant FDR-corrected within-condition phase consistencies are indicated by gray * markers; significant FDR-corrected between condition phase consistencies are denoted by black * markers across the top of the plot.

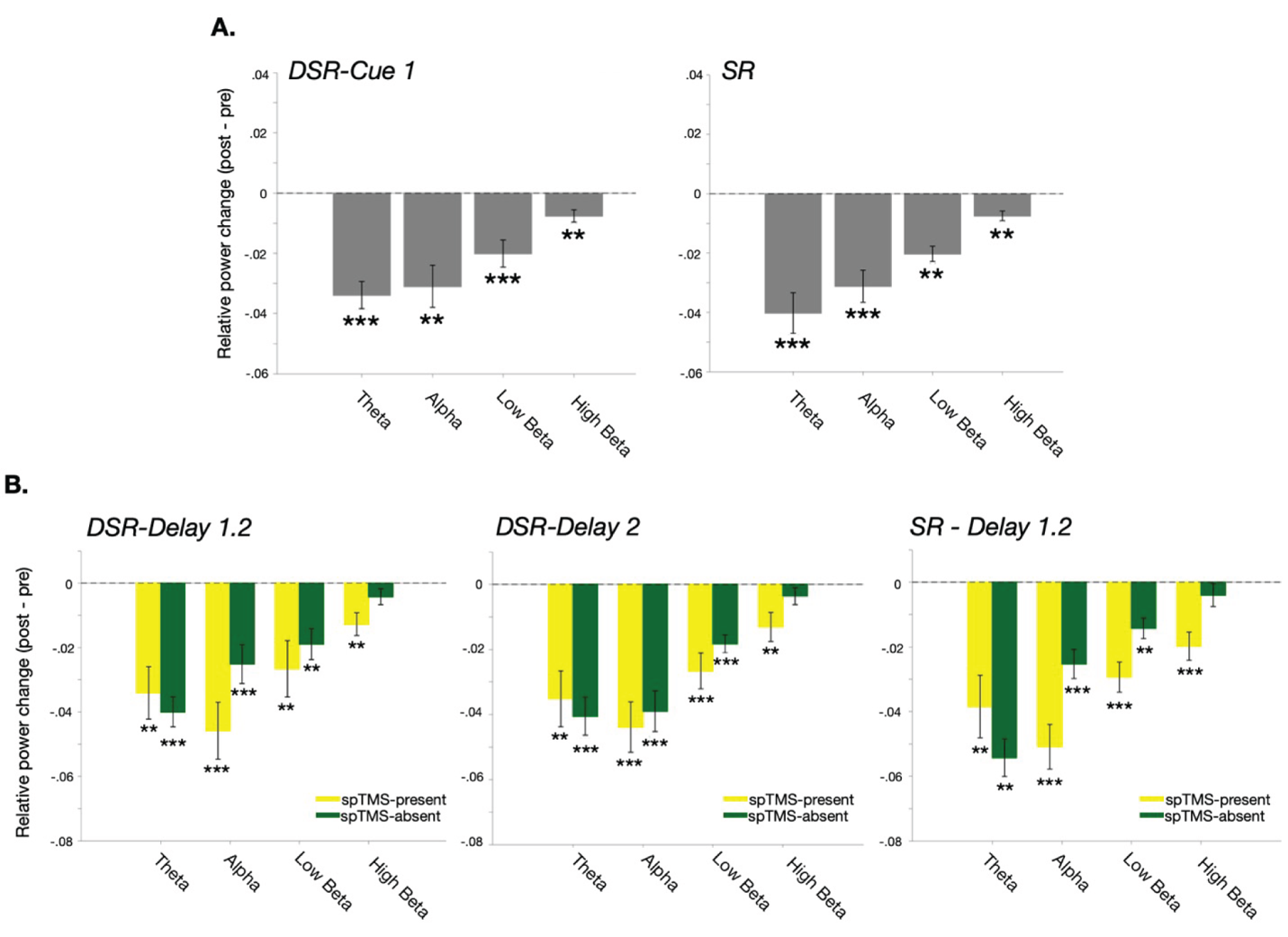

**Figure 6-3. Within-subject power. A.** Cue-evoked changes in relative power (post – pre) for each of the four frequency bands in response to *Cue 1* (DSR; left) and the single cue in the SR task (right). **B.** Same as **A.** for spTMS-evoked changes of relative power, with comparison to baseline in trials when spTMS was not delivered. *** markers: *p* < .001; ** markers: *p* < .01; * markers: *p < .*05 based on permutation tests; error bars correspond to +/- 1 *SEM.*
